# miR-183/96/182 cluster is an important morphogenetic factor targeting *PAX6* expression in differentiating human retinal organoids

**DOI:** 10.1101/2020.04.01.019539

**Authors:** Lucie Peskova, Denisa Jurcikova, Tereza Vanova, Jan Krivanek, Michaela Capandova, Zuzana Sramkova, Magdalena Kolouskova, Hana Kotasova, Libor Streit, Tomas Barta

## Abstract

MicroRNAs (miRNAs), a class of small, non-coding RNA molecules represent important regulators of gene expression. Recent reports have implicated their role in the cell specification process acting as “fine-tuners” to ensure the precise gene expression at the specific stage of cell differentiation. Here we used retinal organoids differentiated from human pluripotent stem cells (hPSCs) as a model to closely investigate the role of a sensory organ-specific and evolutionary conserved miR-183/96/182 cluster. Using a miRNA tough decoy approach, we inhibited the miR-183/96/182 cluster in hPSCs. Inhibition of the miRNA cluster resulted in an increased expansion of neuroepithelium leading to abnormal “bulged” neural retina in organoids, associated with upregulation of neural-specific and retinal-specific genes. Importantly, we identified *PAX6*, a well-known essential gene in neuroectoderm specification, as a target of the miR-183/96/182 cluster members. Taken together, the miR-183/96/182 cluster not only represents an important regulator of *PAX6* expression, but it also plays a crucial role in retinal tissue morphogenesis.

## Introduction

Different classes of non-coding RNAs are dynamically expressed during the development in a tissue-specific manner, and they control diverse biological processes. MiRNAs, a class of small, noncoding RNAs, have emerged as the critical regulatory components of the timing of stepwise events during development, typically by acting as modulators of molecular circuitries. Expression of miRNAs appears to be highly influenced by the developmental stage and tissue specificity representing the key regulatory elements of virtually all cell processes underlying cell function and identity.

The miR-183/96/182 cluster comprising of three miRNA members (miR-183, miR-182 and miR-96) is a highly conserved miRNA cluster in bilaterian organisms. All miRNA members of this cluster share high sequence homology and are found to be specifically expressed in pluripotent stem cells, the retina and other sensory organs as well (reviewed in [1]). Given its target genes, evolutionary conservation and tissue-specific expression pattern in sensory organs and pluripotent stem cells, the miR-183/96/182 cluster is likely an important factor during the differentiation of pluripotent stem cells into neural tissues and sensory organs.

Here we aimed to elucidate the role of the miR-183/96/182 cluster in the differentiation process of human embryonic stem cells (hESCs) and induced pluripotent stem cells (hiPSCs) into retinal organoids, well-established *in vitro* models recapitulating spatial and temporal differentiation of the retina. Inhibition of the miR-183/96/182 cluster, using a though decoy (TuD) approach, had a profound effect on the organoid morphology with evident expansion of the neuroepithelium leading to an abnormal “bulged” neural retina phenotype, associated with upregulation of neuro- and retina-specific genes. Moreover, we described *PAX6*, a master regulator of neuroectoderm specification and eye development, as a target of the miR-183/96/182 cluster in the differentiating organoids. This work provides a novel insight into the mechanisms of retinal tissue differentiation and morphogenesis and highlights the importance of the miR-183/96/182 cluster in these processes.

## Methods

### Cell culture and hiPSCs generation

Skin fibroblasts from healthy, 8-years-old male donor were reprogrammed using an Epi5^™^ Episomal iPSC Reprogramming Kit (Invitrogen) according to the manufacturer’s instructions. hiPSCs were maintained in Essential 8 culture media (Gibco) on Vitronectin-coated dishes and regularly passaged every ~4 days. hiPSCs between passages 30-40 were used in this study (Supplementary Figure 1). hESCs (H9 cell line - herein referred to as hESCs) was obtained from the WiCell Bank (Wisconsin) and was cultivated using the same conditions, as described for hiPSCs.

### TuD design, generation and cloning

Clustered TuD was designed and generated, as described in [2]. Briefly, binding sites (BS) for miR-96-5p, miR-182-5p, miR-183-5p, and a scrambled sequence (SCR) that does not target any miRNA of this cluster (or similar miRNA) were designed and *in silico* tested using the miRNAsong tool [3]. BS were designed to contain mismatches, which upon binding to the miRNA, create a bulge that prevents cleavage by Argonaute RISC Catalytic Component 2 (Ago2) and improve the stability of TuD transcript [2] (TuD and SCR sequences are shown in Supplementary Figure 2). The structure of clustered TuD, containing binding sites for miR-96-5p, miR-182-5p, miR-183-5p flanked with AgeI and EcoRI restriction sites, was analysed using RNAfold (http://rna.tbi.univie.ac.at/cgi-bin/RNAWebSuite/RNAfold.cgi) (Supplementary Figure 3). TuD was synthetized as a minigene (Integrated DNA Technologies) and cloned to a lentiviral vector containing DOX-inducible U6 promoter and TetRep-P2A-Puro-P2A-mCherry [4] (vector kindly provided by Mikael Altun). Lentiviral particles were generated as described previously [3,5] using pMD2.G (Addgene #12259) and psPAX2 (Addgene #12260) (gift from Didier Trono, École Polytechnique Fédérale de Lausanne, Lausanne, Switzerland). After transduction, mCherry positive hiPSCs or hESCs were sorted using BD FACSAria^™^ II (BD Biosciences).

### PAX6 3 ‘UTR reporter

Lentiviral vector expressing destabilized EGFP (d2EGFP) followed by *PAX6* 3’UTR sequence was designed and constructed using VectorBuilder (https://en.vectorbuilder.com/) (Figure 6B). Lentiviral particles were generated as described above. Cell colonies expressing d2EGFP were observed using a Cell^R imaging station (Olympus) at 1 second exposure time.

### Ago2-immunoprecipatation (IP) and PCR

Ago2-IP was performed as described previously[3]. Briefly, cells were washed three times with phosphate-buffered saline (PBS) (pH 7.4) and UVC-irradiated (0.2 mW/cm^2^ for 60 seconds). Cells were then lysed in a buffer containing 50 mM Tris-HCl (pH 7.4), 150 mM NaCl, 5 mM MgCl2, 0.5% NP-40, 15 mM ethylenediaminetetraacetic acid (EDTA), freshly added complete mini EDTA-free protease inhibitor cocktail (Roche), and RNAse inhibitor (final concentration 2 U/ml) (Applied Biosystems^™^). Cell lysates were cleared by centrifugation at 15,000 × *g* for 10 minutes at 4 °C, the supernatant was pre-cleared by incubation with protein G sepharose beads (GE Healthcare) on a rotating platform for 1 hour at 4°C, and the sepharose beads were centrifuged. The supernatant was then incubated on ice for 1 hour with the primary antibody EIF2C2 (H00027161-M01, Abnova). Immunocomplexes were collected overnight at 4°C on protein G sepharose beads and washed five times with a lysis buffer. In order to detect Ago2, immunocomplexes were subjected to western blot analysis using the Ago2 primary antibody (#2897, Cell Signalling Technology). RT-qPCR was performed with RNA isolated using Trizol reagent and reverse transcribed using a High-Capacity cDNA Reverse Transcription Kit (Applied Biosystems) with random hexamer primers. The RT product was amplified by real-time PCR (Roche LightCycler^®^ 480 PCR instrument) using PowerUp^™^ SYBR^™^ Green Master Mix (Applied Biosystems^™^). The following set of primers was used for detection of TuD miR-183/96/182 expression: forward 5’-GTGCCAAACTTCGTGAAGGG, reverse 5’-TGTGAATTGTGATGGCGCTC. In total four reactions were performed: Ago2-IP (SCR, DOX-), Ago2-IP (SCR, DOX+), Ago2-IP (TuD miR-183/96/182, DOX-), and Ago2-IP (TuD miR-183/96/182, DOX+). The TuD miR-183/96/182 enrichment in Ago2-IP (TuD miR-183/96/182, DOX+) was normalised to the Ago2-IP (TuD miR-183/96/182, DOX-), as the expression of TuD miR-183/96/182 was not detected in the SCR control.

### Retinal organoids generation

Retinal organoids were generated from both, hiPSCs as well as hESCs, according to the protocol described elsewhere [6,7] with slight modifications (Figure 3A). Briefly, cells in E8 medium were seeded into a V-shaped 96-well plate (6 000 cells/well) coated with Polyhema (Merck). After 48 hours (Day 0 of the differentiation process) the culture medium was changed to gfCDM [6]. On day 6 recombinant human BMP4 (Peprotech) was added to the culture to the final concentration 2.2 nM and then the medium was changed every 3rd day. On day 18 of the differentiation process, gfCDM was changed to a NR medium [6]. The organoids were cultured in 96 well plates and analysed between days 18-35.

### Scanning electron microscopy (SEM)

The organoids were fixed with 3% glutaraldehyde in a 0.1 M cacodylate buffer, then washed three times with 0.1 M cacodylate buffer, dehydrated using ascending ethanol grade (30, 50, 70, 80, 90, 96, and 100%) and dried in a critical point dryer (CPD 030, BAL-TEC Inc., Liechtenstein) using liquid carbon dioxide. The dried organoids were sputtered with gold in a sputter coater (SCD 040, Balzers Union Limited) and observed in a scanning electron microscope (VEGA TS 5136 XM, Tescan Orsay Holding, Czech Republic) using a secondary emission detector and a 20 kV acceleration voltage.

### Flow Cytometry

Cells were harvested using 0.5 mM EDTA in PBS, washed with PBS, and then fixed with ice-cold 70% ethanol. The fixed cells were washed twice with wash buffer (5mM EDTA, 2% Foetal Bovine Serum in PBS) and incubated for 1 hour with the primary antibodies SOX2 (NB110-37235, Novus Biological, 1:400) or Oct-3/4 (sc-5279, Santa Cruz, 1:400). The cells were then washed and incubated with secondary antibodies antiRabbit or anti-Mouse AlexaFluor 488 (Thermo Fisher Scientific, 1:1000) for 1 hour. For TRA-1-60 expression assessment, non-fixed cells and the TRA-1-60 antibody (MAB4360, Millipore, 1:100) were used. Flow cytometry analysis was performed using BD FACSAria^™^ (BD Biosciences). Datasets were analysed using FlowJo software (www.flowjo.com).

### Western blot analysis

Western blot analysis was performed, as described elsewhere [8]. Briefly, at least 16 pooled organoids per each condition were washed three times with PBS (pH 7.4) and lysed in buffer containing 50 mM Tris-HCl (pH 6.8), 10% glycerol and 1% sodium dodecyl sulfate (SDS). The lysates were homogenized by sonication and protein concentrations were determined using DC Protein Assay (Bio-Rad). The lysates were then supplemented with 0.01% bromophenol blue and 1% β-mercaptoethanol and denatured at 100 °C for 5 minutes. The prepared samples were separated by SDS-polyacrylamide gel electrophoresis and transferred onto a polyvinylidene fluoride (PVDF) membrane (Merck Millipore). The PVDF membrane was blocked in 5% skimmed milk in Tris-buffered saline containing Tween for 1 hour and incubated with primary antibodies overnight at 4 °C. The following antibodies were used: PAX6 (sc-53108, Santa Cruz, 1:1000), SOX2 (sc-365823, Santa Cruz, 1:1000), SOX1 (bs-12276R, Bioss, 1:1000), β-ACTIN (#4970S, Cell Signaling Technology, 1:1000).

### RT-qPCR analysis

At least 12 pooled retinal organoids per sample were homogenised using a 1 ml insulin syringe in 1 ml RNA Blue Reagent (an analogue of Trizol) (Top-Bio), followed by chloroform RNA extraction. RNA was then reverse transcribed using a High-Capacity cDNA Reverse Transcription Kit (Applied Biosystems^™^). The RT product was amplified by real-time PCR (Roche LightCycler^®^ 480 PCR instrument) using PowerUp^™^ SYBR^™^ Green Master Mix (Applied Biosystems^™^). Primer sequences are shown in Supplementary Figure 4. For the determination of miRNA expression, RNA was reverse transcribed (16 °C, 30 minutes; 42 °C, 30 minutes; 85 °C, 5 minutes) using a TaqMan^®^ MicroRNA Reverse Transcription Kit and specific primers for RNU6B, miR-96-5p, miR-182-5p, and miR-183-5p (Applied Biosystems). Reverse transcription products were then amplified by quantitative PCR (95 °C, 5 minutes; 95 °C, 15 seconds; 60 °C, 60 seconds; 40 cycles) (LightCycler^®^ 480, Roche) using TaqMan Universal PCR Master Mix and specific probes for miRNAs. Relative microRNA expression was determined using the ΔΔCt method and normalized to endogenous control RNU6B.

### miR-183/96/182 targets network analysis

A list of all experimentally validated target genes of the selected miRNAs was downloaded from miRTarBase (http://mirtarbase.mbc.nctu.edu.tw/) [9], the list of target genes was then analysed using the DAVID tool (https://david.ncifcrf.gov/) [10] and visualised using Cytoscape v3.7.2 [11,12]. Please note that only genes that are targeted by at least two members of the miR-183/96/182 cluster were processed.

### Analysis of organoid circularity

96 well plates containing organoids were scanned using an Epson perfection V550 scanner. To remove potential bias for the bulged or normal phenotype, images were analysed *in silico* using the ImageJ – Analyze particles tool, with circularity threshold set to 0.6-1. Organoids that did not pass this threshold were considered as abnormal (“bulged”) (Supplementary Figure 5).

### Whole-mount analysis of organoids

Clearing and staining protocol was adapted from [13,14]. Briefly, the organoids were fixed in 4% paraformaldehyde for 30 minutes at room temperature, incubated in CUBIC-1 solution for 24 hours at 37 °C, washed in washing solution (PBS with 0.1% Tween20 (P1379, Sigma Aldrich) and 0.5% Triton X-100 (T8787, Sigma Aldrich)), incubated with primary antibodies: ZO1 (Abcam, ab190085, 1:100) and SOX1 (Bioss, bs-12276R, 1:200) with DAPI (D9542, Sigma Aldrich) dissolved in washing solution for 24 hours at room temperature, washed in washing solution, incubated with appropriate Alexa Fluor (Thermo Fisher Scientific) secondary antibodies dissolved in washing solution (anti-Goat 488 (1:500, A-11055) and anti-Rabbit 568 (1:500, A-10042)) for 4 hours at room temperature and finally cleared by CUBIC-2 solution overnight at room temperature. The cleared and stained organoids were embedded in 3% low-gelling agarose (A9414, Sigma Aldrich) dissolved in a mixture of CUBIC-2 and PBS (3:1) on the bottom of Ibidi μ-Slide 8 well dishes (80826, Ibidi) and imaged on a confocal microscope ZEISS LSM 880 using 10x objective. The raw files were processed using IMARIS software (Bitplane, Zürich, Switzerland) and the final images and movies were exported.

### Immunohistochemistry

The organoids were fixed in 4% paraformaldehyde for 30 minutes at room temperature, washed with PBS, cryopreserved in 30% sucrose (16104, Sigma Aldrich) in PBS for 1 hour at room temperature, embedded in Tissue-Tek^®^ O.C.T. Compound medium (Sakura) and sectioned on a cryostat (CM1850, Leica), (~10 μm). The sections were washed with PBS and blocked in Blocking buffer (PBS with 0.3% Triton X-100 (T8787, Sigma Aldrich) and 5% normal goat serum (G9023, Sigma Aldrich)) for 1 hour at room temperature in a humidified chamber. The primary antibodies were diluted in Antibody diluent (PBS with 0.3% Triton X-100 and 1% bovine serum albumin (A9647, Sigma Aldrich)) and applied to the sections overnight at 4 °C in a humidified chamber. The following primary antibodies were used: PAX6 (sc-53108, Santa Cruz, 1:200) and RAX (sc-271889, Santa Cruz, 1:200). The sections were washed with Antibody diluent and a secondary antibody (anti-Mouse AlexaFluor 488 (Thermo Fisher Scientific, 1:1000)) was applied for 1 hour at room temperature in a humidified chamber. The nuclei were stained with 1 μg/ml DAPI in PBS for 4 minutes at room temperature and the sections were mounted using Fluoromount^™^ Aqueous Mounting Medium (F4680, Sigma Aldrich).

## Results

### miR-183/96/182 cluster expression during differentiation of hiPSCs into retinal organoids

The polycistronic miR-183/96/182 cluster is an evolutionary conserved miRNA cluster containing three highly similar miRNA members. Despite the sequence homology, and hence the crosstalk between individual members of the cluster, there are also small differences in their sequence leading to different mRNA targets of individual members of the cluster, however often within the same functional group expressed in the same tissues [1]. Our network analysis shows that the individual members share multiple targets with most of them being expressed in the brain or the eye, corroborating previous findings that the miR-183/96/182 cluster is expressed and active in sensory organs, and neural tissues (Figure 1A).

**Figure 1:**
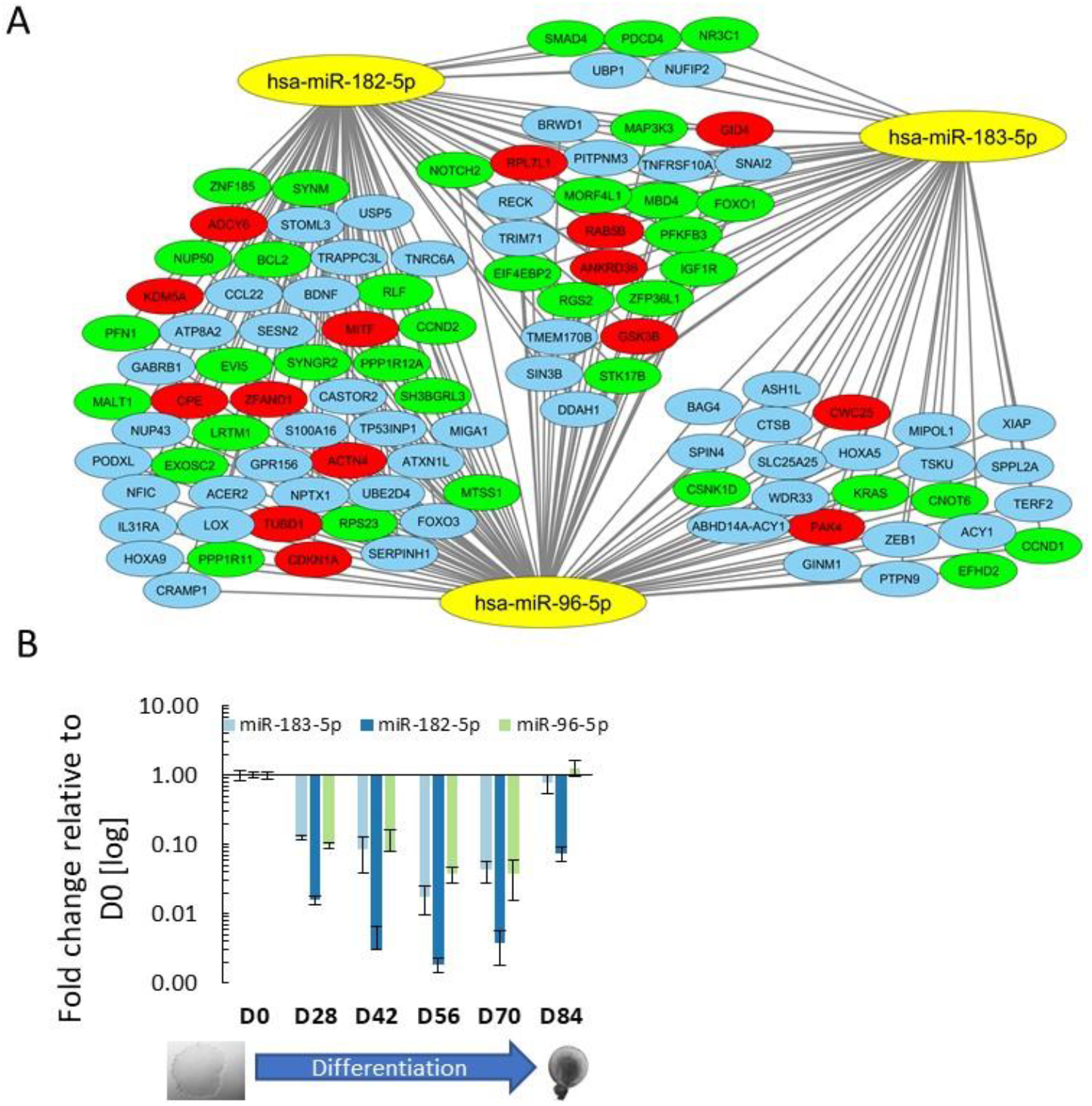
**A)** A graphical representation of the genes targeted by the members of the miR-183/96/182 cluster. Only experimentally validated genes that are targeted by at least two members of the cluster are shown. Genes known to be expressed in the brain or eye are highlighted in green or red respectively. **B)** Expression of individual members of the miR-183/96/182 cluster during differentiation of hiPSCs to the retinal organoids, as determined by RT-qPCR.

We used retinal organoids as a model to closely investigate the role of the cluster in the differentiation process toward neural and sensory tissues. We first sought to determine the expression of the miR-183/96/182 cluster during differentiation of hiPSCs into retinal organoids. Members of the miR-183/96/182 cluster are expressed in hiPSCs and their expression is gradually decreased over time to day 70 (D70), however the expression is then upregulated at D84 to the similar level, as detected in pluripotent hiPSCs (Figure 1B). This expression pattern suggests that the miR-183/96/182 plays some role during the early differentiation steps of pluripotent stem cells into retinal organoids.

### Generation and testing of TuD for the inhibition of the miR-183/96/182 cluster

To study the role of the miR-183/96/182 cluster in the differentiation process of hPSCs into retinal organoids, we aimed to perform loss-of-function experiments. To inhibit the miR-183/96/182 cluster, we designed a miRNA TuD [2] containing binding sites for the individual members of the cluster and cloned it into Doxycycline(DOX)-inducible lentiviral vector (Figure 2A, S2A, B, and S3). Upon transduction and antibiotic selection, mCherry positive cells were sorted. Generated stabile cell lines maintained mCherry expression for at least 15 passages (Fig 2B, C).

**Figure 2:**
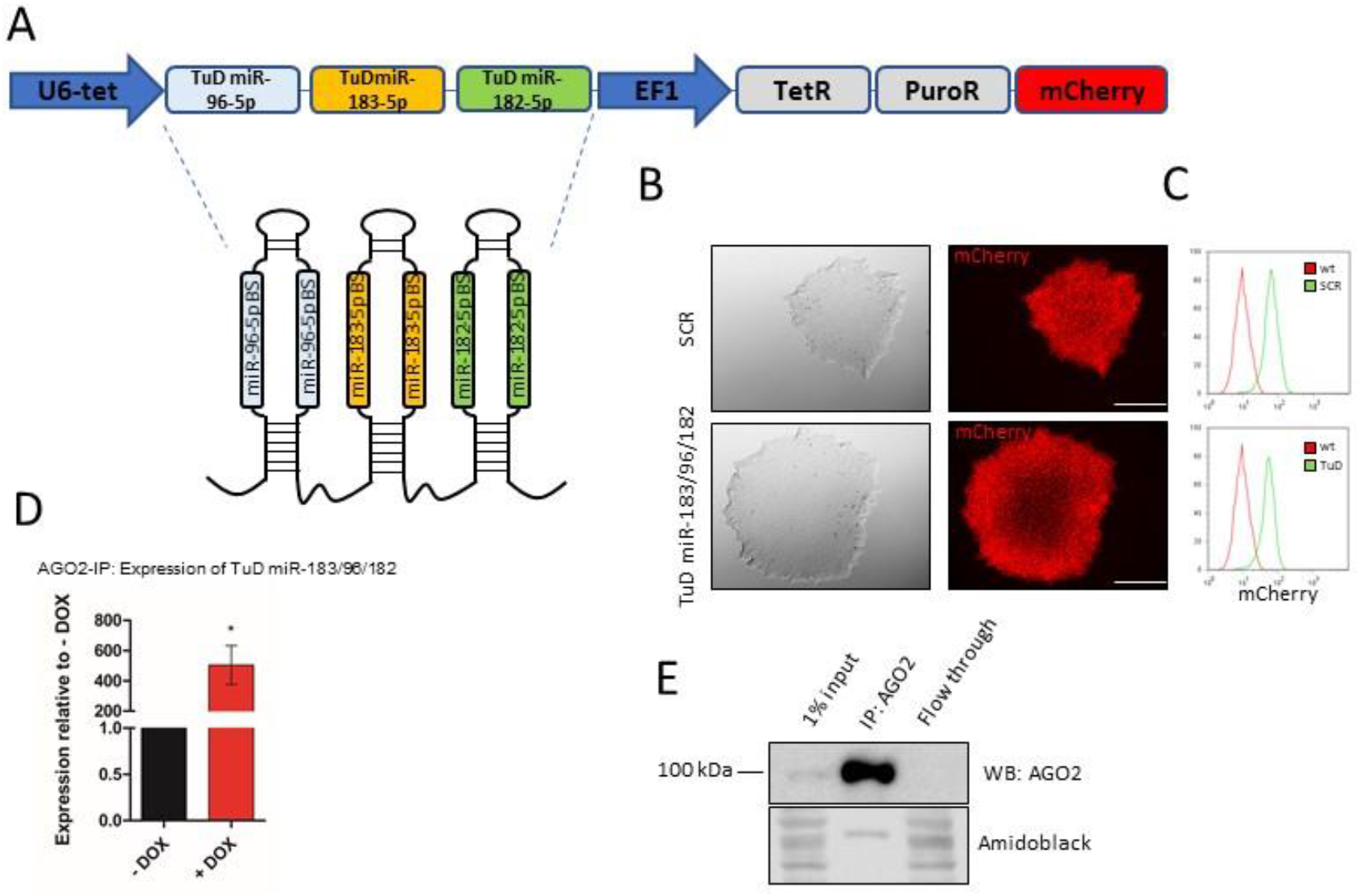
**A)** Schematic presentation of the vector composition for miR-183/96/182 inducible inhibition using the TuD approach. **B)** The morphology of transduced hiPSCs expressing T uD for inhibition of the miR-183/96/182 cluster or SCR control, as determined by bright field and fluorescent microscopy. **C)** Quantification of cells expressing mCherry, as determined by flow cytometry. **D)** Quantification of TuD miR-183/96/182 transcript levels in Ago2-immunoprecipitated (Ago2-IP) fraction, as demonstrated by RT-qPCR. Data are depicted as mean + SD of triplicates. **E)** Western blot analysis of Ago2 levels in Ago2-IP reaction. Lower part: amidoblack-stained PVDF membrane represents a loading control.

**Figure 3:**
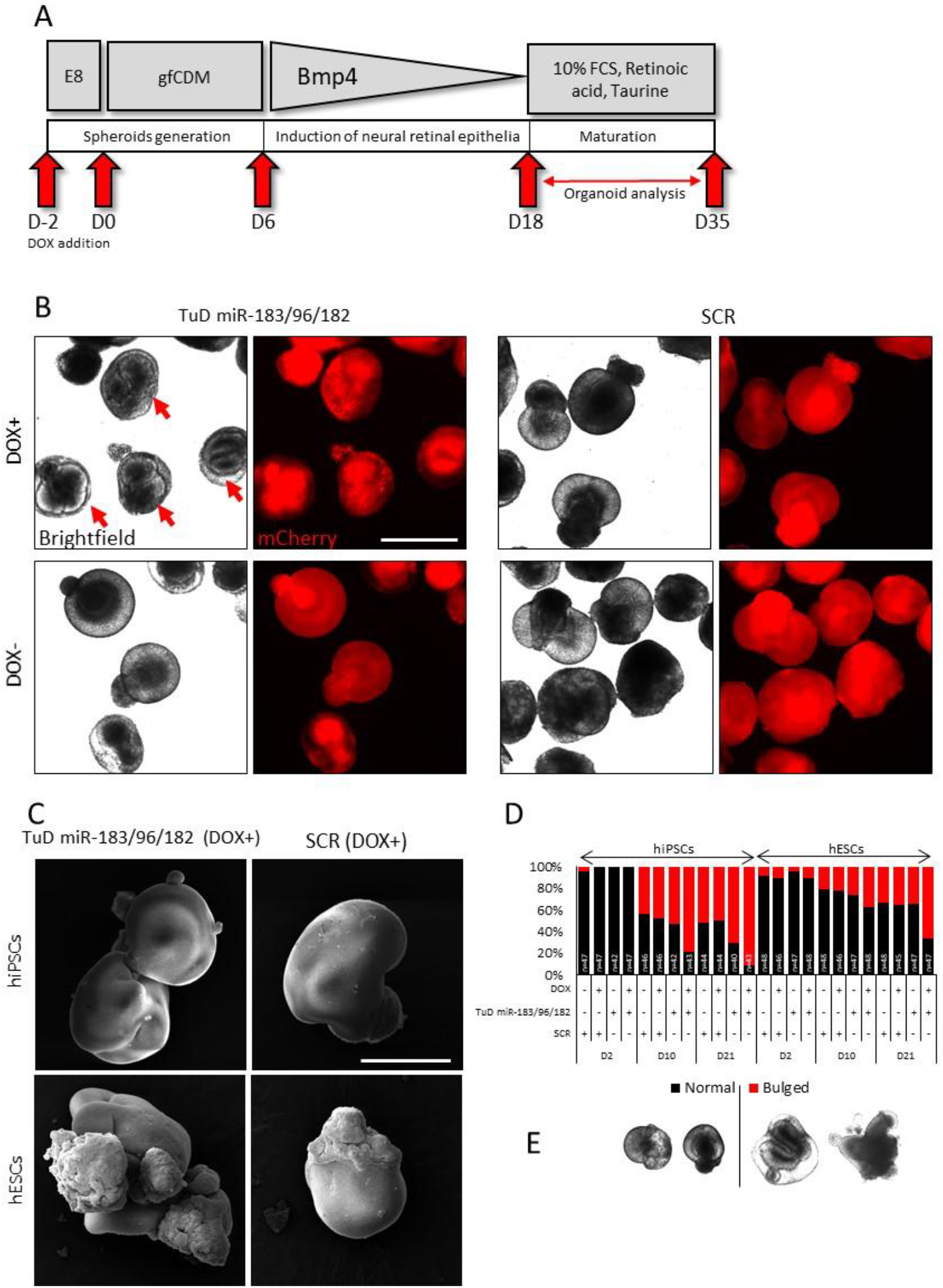
**A)** Schematic presentation of the differentiation protocol. **B)** Morphology of the retinal organoids at D18 of differentiation, as determined by bright field and fluorescent microscopy. Arrows point to abnormal, bulged retinal organoids. Scale bar = 2 mm. **C)** Morphology of retinal organoids at D35 of differentiation, as determined by SEM. Scale bar = 500 μm. **D)** Quantification of bulged organoids. **E)** Representative images of normal retinal organoids that passed the circularity threshold during *in silico* circularity analysis and bulged organoids that did not pass the circularity threshold.

To prove the functionality of the TuD miR-183/96/182, we first determined, if the miRNA-containing RNA-induced silencing complex (RISC) was associated with the TuD miR-183/96/182 transcript. We tested if the TuD miR-183/96/182 transcripts were enriched in immunoprecipitated Ago2. RT-qPCR revealed that the TuD transcripts were strongly enriched in the Ago2-IP (TuD miR-183/96/182 DOX+) fraction compared to the Ago2-IP (TuD miR-183/96/182 DOX-) control (Figure 2D, E), indicating that the TuD transcripts were associated with the Ago2-containing RISC complex. As an additional control, we used cells expressing an SCR sequence, however we could not detect the TuD miR-183/96/182 transcript in either of the Ago2-IP (SCR DOX+/-) fractions (data not shown).

### Inhibition of the miR-183/96/182 during retinal organoid differentiation induces neuroepithelial expansion

Given the experimentally validated mRNA targets of the miR-183/96/182 cluster and its potential role in the differentiation towards neuroepithelium and sensory tissues, we hypothesized that inhibition of the cluster will lead to increased efficiency of differentiation. To test this, we inhibited the miR-183/96/182 cluster, using DOX-induced expression of TuD miR-183/96/182, two days prior to the onset of hPSCs differentiation (Day-2) (DOX concentration = 1 μg/ml) (Figure 3A). The morphology of retinal organoids at the early stage of differentiation (between D17-D35) was analysed using bright field, fluorescent, and scanning electron microscopy. Inhibition of the miR-183/96/182 cluster resulted in the expansion of neuroepithelium in the retinal organoids derived from both hiPSCs as well as hESCs, visible as extra bulged structures in the organoids leading to the generation of abnormal, bulged retinal organoids (Figure 3B, C).

It is of note that abnormal organoids normally appear in a culture in low numbers, however when the miR-183/82/182 cluster is inhibited their number greatly increases. In order to assess the penetration of the bulged phenotype we analysed the roundness of retinal organoids. Using an unbiased *in silico* approach (see Methods section), we analysed the morphology of the generated retinal organoids at D2, D10, D21 and quantified the numbers of bulged organoids generated from hiPSCs and hESCs. Bulged organoids were visible as early as at D10 of the differentiation process, when the neuroepithelium typically appears [6]. Inhibition of the miR-183/96/182 cluster led to the generation of ~80% (at D10) and ~90% (at D25) of bulged organoids generated from hiPSCs, while ~40% (at D10) and ~65% (at D25) of bulged organoids were generated from hESCs (Figure 3D, E, Supplementary Figure 5).

### Inhibition of the miR-183/96/182 cluster leads to upregulation of neural and retinal markers in differentiating retinal organoids

Inhibition of the miR-183/96/182 cluster leads to the generation of abnormal retinal organoids. We next assessed whether the abnormal morphology is associated with an upregulated expression of retina-specific genes. Western blot and RT-qPCR analysis revealed upregulated expression of retina-specific genes *RAX, VSX2* and neuroectoderm-specific genes *SOX1, SOX2*, and *PAX6* in both hiPSCs-as well as hESCs-derived retinal organoids (Figure 4A, B, C, D). RT-qPCR analysis also revealed that the upregulation of these genes is slightly higher in hiPSCs-derived organoids (~2-3.5 fold change) when compared to hESCs-derived organoids (~1.5-2 fold change) (Figure 4B, D). This is in line with the observed lower numbers of bulged organoids derived from hESCs (Figure 3D).

**Figure 4:**
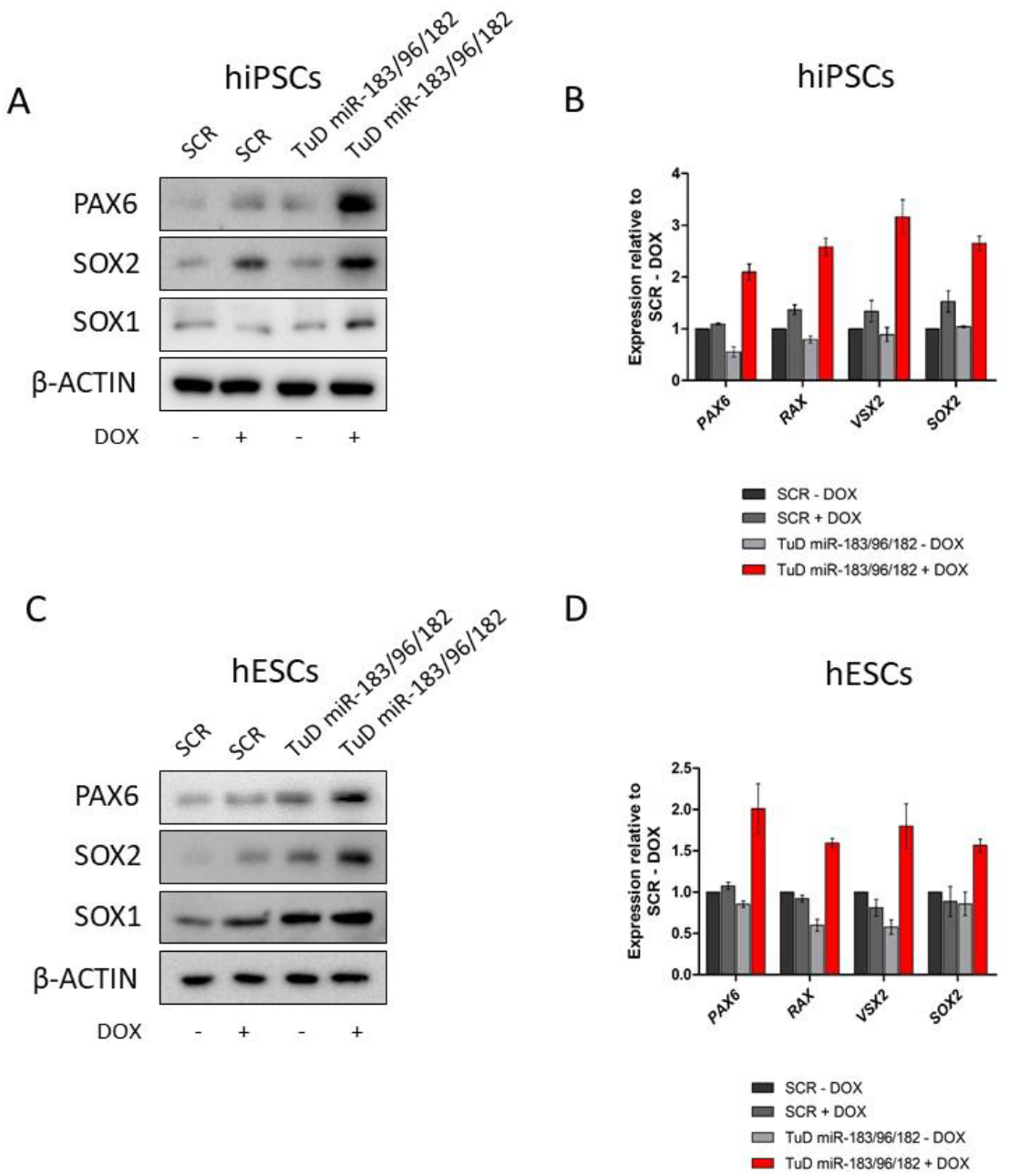
**A)** Expression of PAX6, SOX2, and SOX1 in the retinal organoids (at D25) derived from hiPSCs, as determined by western blot analysis. β-ACTIN represents a loading control. **B)** Expression of *PAX6, RAX, VSX2*, and *SOX2* in the retinal organoids (at D25) derived from hiPSCs, as determined by RT-qPCR analysis. Relative gene expression was determined using the ΔΔCt method and normalized to endogenous control *GAPDH*. **C)** Expression of PAX6, SOX2, and SOX1 in the retinal organoids (at D25) derived from hESCs, as determined by western blot analysis. β-ACTIN represents a loading control. **D)** Expression of *PAX6, RAX, VSX2*, and *SOX2* in the retinal organoids (at D25) derived from hESCs, as determined by RT-qPCR analysis. Relative gene expression was determined using the ΔΔCt method and normalized to endogenous control *GAPDH*.

**Figure 5:**
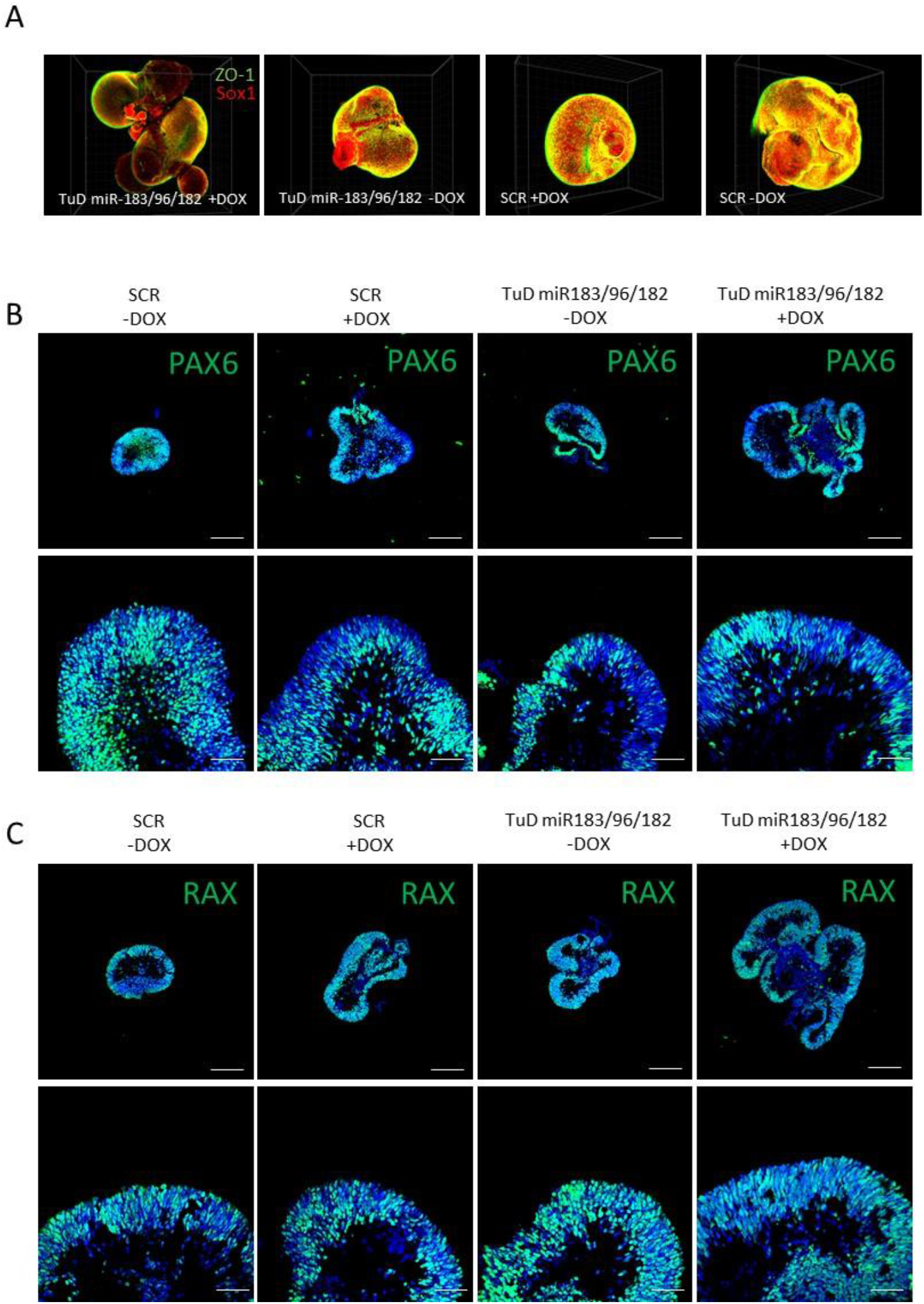
**A)** ZO-1 and Sox1 expression, as determined by whole-mount organoid analysis using confocal microscopy of the organoids at D25. **B, C)** Expression of PAX6 and RAX, as determined by confocal microscopy of cryo-sectioned organoids (at D25) along the central-peripheral axis. Scale bar on the upper panels = 500 μm. Scale bar on the bottom panels = 80 μm

### Expanded neuroepithelium is positive for neuroectodermal and retinal markers

Given the abnormal phenotype of the retinal organoids upon miR-183/96/182 inhibition, we sought to analyse whether the abnormal structures represent early neural retina. We performed immunohistochemistry of cryo-sectioned retinal organoids and whole-mount organoid analysis. The expanded structures were positive for SOX1, PAX6, RAX, and ZO-1 markers typically found in the neuroectoderm and early neural retina[6]. Taken together, our results indicate that abnormal retinal organoids contain expanded neural retina structures, suggesting that the inhibition of the miR-183/96/182 cluster leads to enhanced differentiation of hPSCs into neural retina.

### The miR-183/96/182 cluster targets PAX6

By screening for additional genes (using the miRmap tool, https://mirmap.ezlab.org/) [15] that are potentially regulated by the miR-183/96/182 cluster, we identified *PAX6* as a predicted target of all members of the cluster (Figure 6A). We identified two putative binding sites on the *PAX6* 3’UTR sequence. One binding site is located at the 3’UTR 981 bp and is shared by miR-96-5p and miR-182-5p, the other site potentially binds miR-183-5p and is located at the 3’UTR 3236 bp. Therefore, we aimed to assess, whether the miRNA cluster targets *PAX6* mRNA. We infected hiPSCs expressing DOX-inducible TuD miR-183/96/182 with lentiviral particles containing vector expressing destabilized GFP (d2EGFP) followed by the 3’UTR *PAX6* sequence (Figure 6B). We detected a very dim GFP fluorescent signal in the presence of DOX, indicating that *PAX6* mRNA is targeted by the miR-183/96/182 cluster by binding to the *PAX6* 3’UTR sequence (Figure 6C).

**Figure 6:**
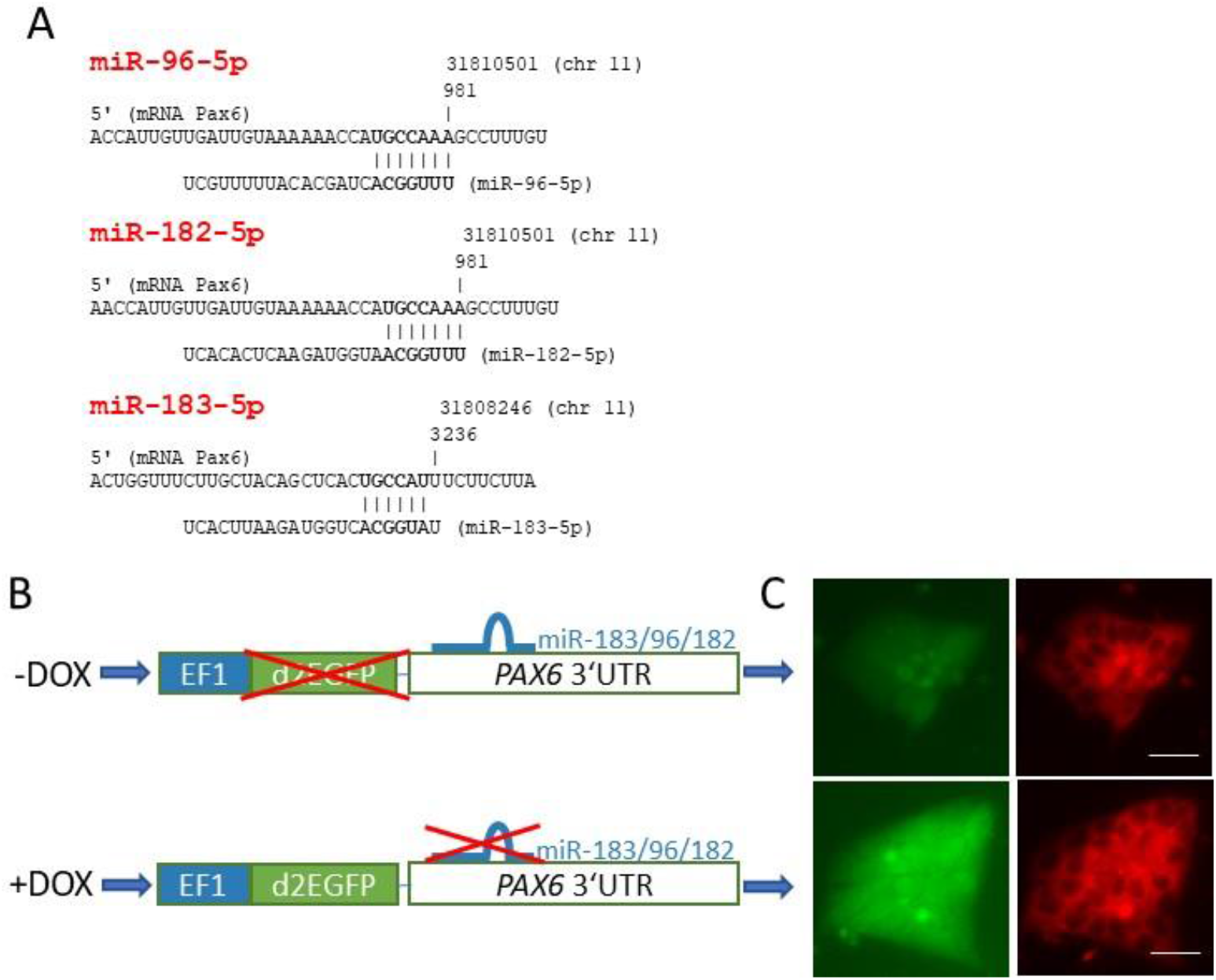
**A)** Target sites of the individual members of the miR-183/96/182 cluster on *PAX6* mRNA, as predicted by the miRmap tool. **B)** Schematic presentation of *PAX6* 3’UTR fluorescent reporter in the presence or absence of Doxycycline. **C)** Fluorescence signal from *PAX6* 3’UTR reporter, as determined by fluorescent microscopy. Pictures were taken using the same exposure time (1 second) and the same microscope settings. Scale bar = 40 μm.

## Discussion

MiRNAs represent the key “fine-tuners” of gene expression during the differentiation of hPSCs. It has been previously shown that manipulation with single miRNA or the whole miRNA cluster/family may have a significant impact on differentiation outcomes of pluripotent stem cells [16-19]. Here we show that the miR-183/96/182 cluster regulates *PAX6* expression in differentiating hPSCs into retinal organoids. Inhibition of the miR-183/96/182 cluster, using the clustered TuD miRNA suppression approach, leads to increased neural retina expansion in retinal organoids associated with upregulated expression of *PAX6* and other neural and retinal-specific genes. Despite the fact that we have shown *PAX6* as a direct target of the miR-183/96/182 cluster, it is still not clear, whether the observed bulged phenotype of retinal organoids is directly mediated by upregulated *PAX6* expression or other miR-183/96/182 target genes. *PAX6* belongs to the group of eye field transcription factors (*PAX6, RAX, SIX3, LHX2* and others) and its expression is necessary for eye formation and retinal development [20]. It has been demonstrated that forced expression of *PAX6* induces formation of ectopic retinal tissues [21-23]. Conversely, a study of *PAX6* overexpression effect on mouse eye formation showed abnormalities and progressive degeneration of the eye and retina from day E14.5, however the retina contained normal range of cell types, including retinal ganglion cells [24]. Therefore, it seems that maintaining the correct levels of *PAX6* expression is of vital importance for normal eye development and miRNAs may play an essential role to ensure the correct *PAX6* expression. Although, the role of the miR-96-*PAX6* axis in hPSCs differentiation has been previously implicated [25], we have for the first time demonstrated that the inhibition of the whole miR-183/96/182 cluster has a profound effect on neural retina expansion and morphology of the retinal organoids, hence bringing a novel insight into possible mechanisms of the delicate regulation of *PAX6* in retinal tissues.

It is of note that the miR-183/96/182 cluster is highly-expressed in pluripotent stem cells and sensory organs including the retina [1]. Our data show that the miR-183/96/182 cluster is gradually down-regulated during the initial steps of hPSCs differentiation into the retinal organoids, reaching minimal expression around D55. Later, around D70, its expression is gradually upregulated to the same level, as detected in hPSCs. Gradual time-dependent upregulation of the miR-183/96/182 cluster has been observed also *in vivo* – the expression of individual members of the cluster in the eye is very low during embryonic development and profoundly increases postnatally [26,27]. Possibly, the miR-183/96/182 cluster expression needs to be kept low during the initial phases of retinal tissue development in order to help maintain the necessary level of *PAX6*. In our retinal organoid system, forced inhibition of the miR-183/96/182 cluster resulted in higher expression of retinal and neural-specific genes *PAX6* and *RAX* associated with neural retina expansion. Later, *PAX6* expression is switched off in a certain population of cells that will differentiate into photoreceptors, and the level of the miR-183/96/182 cluster is upregulated [28].

The critical role of the miR-183/96/182 cluster in the retina has been also demonstrated *in vivo*. Mice knock-outs for the miR-183/96/182 cluster have a normally developed retina with no apparent morphological changes, but when mice lacking the miR-183/96/182 cluster were exposed to light, a rapid degeneration of the retina demonstrated by a significant decrease in thickness of the retinal outer nuclear layer was observed [29]. Contrary to this study, Busskamp et al., 2014, demonstrated that the miR-183/96/182 cluster is the key component for maintenance of mouse outer retinal segments [30]. These findings highlight the importance of the miR-183/96/182 cluster in retina maintenance and function, however they are contradictory to our findings. We have demonstrated that inhibition of the miR-183/96/182 cluster leads to increased neural retina expansion, however the miR-183/96/182 knock-out mice developed an apparently normal retina. Why this fundamental difference exists remains unresolved. However, there are multiple plausible explanations: I) *In vivo* studies used a knock-out approach that forbids the presence of other miRNAs having the same or a very similar seed sequence (miR-34a-5p, miR-34c-5p, miR-1271-5p, miR-8053, to name some) and thus resulting in non-specific or incomplete depletion of miRNA molecules with the same or similar seed sequence. II) There are other, yet unknown, compensatory mechanisms that may exist *in vivo*. III) Retinal organoids are differentiated *in vitro* directly from pluripotent stem cells that cannot simply reflect the complexity of an *in vivo* system. Additionally, our data corroborate with the study by Du et al., 2013. The authors demonstrated that miR-96 family overexpression represses neural specification, while its inhibition promotes neural specification of cells *in vitro* differentiated from hPSCs [25]. In our study, we have focused on the early stages of differentiation of retinal organoids, however the impact of the miR-183/96/182 inhibition on maturated retinal organoids remains yet unresolved and it represents the goal of our further investigation.

## Conclusion

This study shows that the miR-183/96/182 cluster regulates PAX6 expression and its inhibition leads to increased neural retina expansion at early stages of differentiation of human retinal organoids derived from hPSCs. Our results indicate that the miR-183/96/182 cluster plays an important role in morphogenesis of the neural retina.

## Acknowledgement

We thank Klára Koudelková for her excellent technical support. This work has been supported by the projects: Brno Ph.D. Talent, the Faculty of Medicine MU to junior researchers T.B. and L.P., the European Regional Development Fund - Project INBIO (No.CZ.02.1.01/0.0/0.0/16_026/0008451). We acknowledge the core facility CELLIM of CEITEC supported by the Czech-BioImaging large RI project (LM2018129 funded by MEYS CR) for their support with obtaining scientific data presented in this work.

---

**Supplementary Figure 1:**
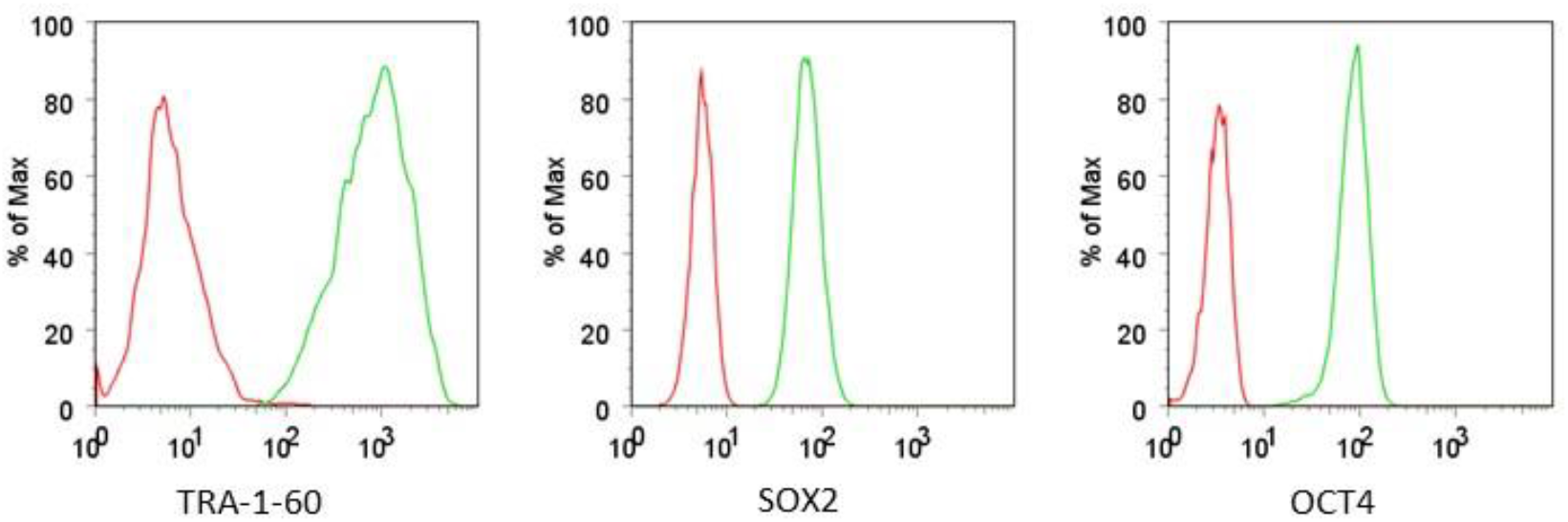
Expression of TRA-1-60, SOX2, and OCT4, as determined by flow cytometry of hiPSCs cells at passage 30.

**Supplementary Figure 2:**
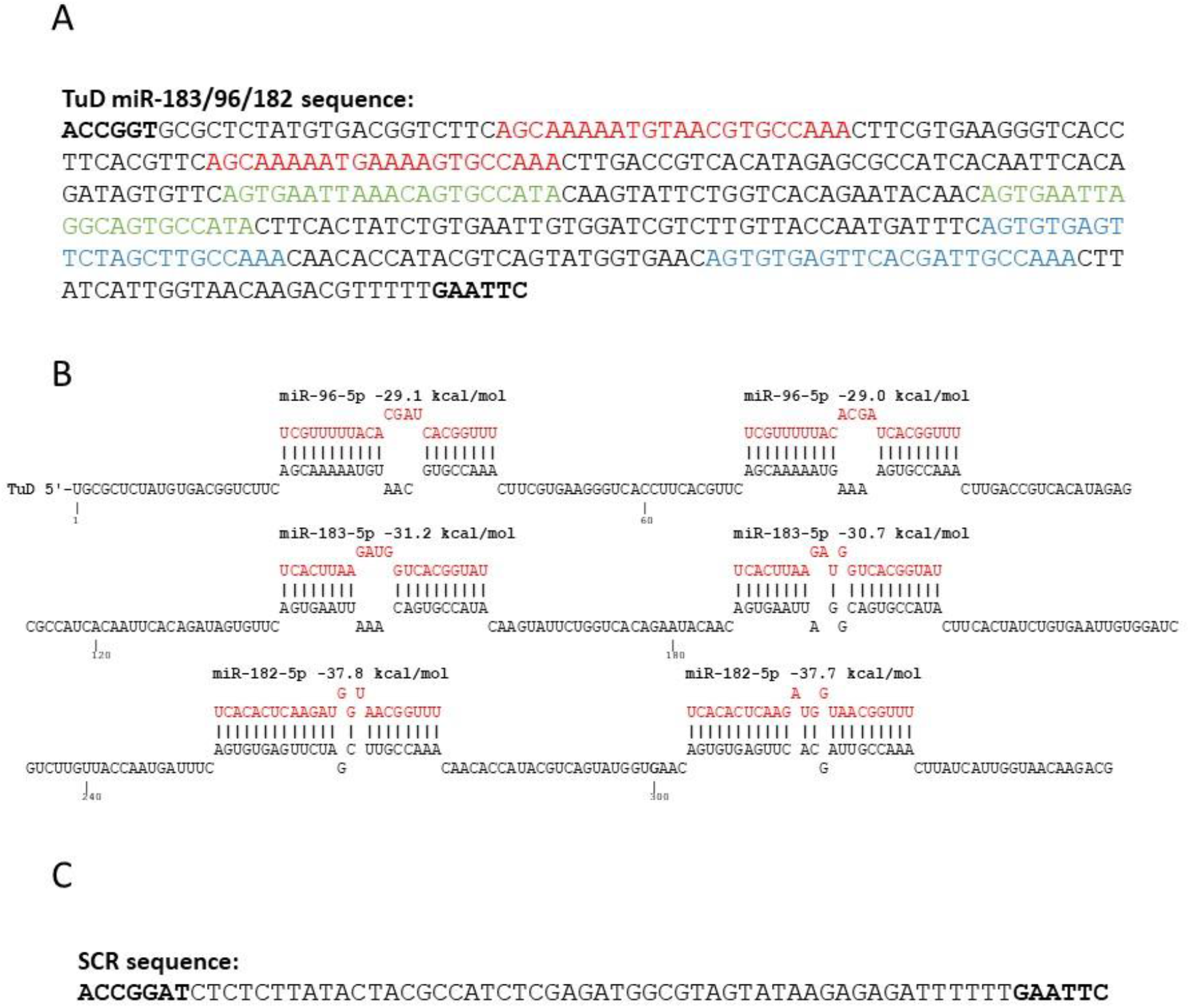
**A)** TuD miR-183/96/182 sequence. Text in red, green, and blue colour represents biding sites for miR-183-5p, miR-96-5p, and miR-182-5p respectively. Bold text represents restriction sites for cloning insert into the vector. **B)** Physical association of individual members of the miR-183/96/182 cluster with TuD transcript. Picture generated by the miRNAsong tool. **C)** SCR sequence. Bold text represents restriction sites for cloning insert into the vector.

**Supplementary Figure 3:**
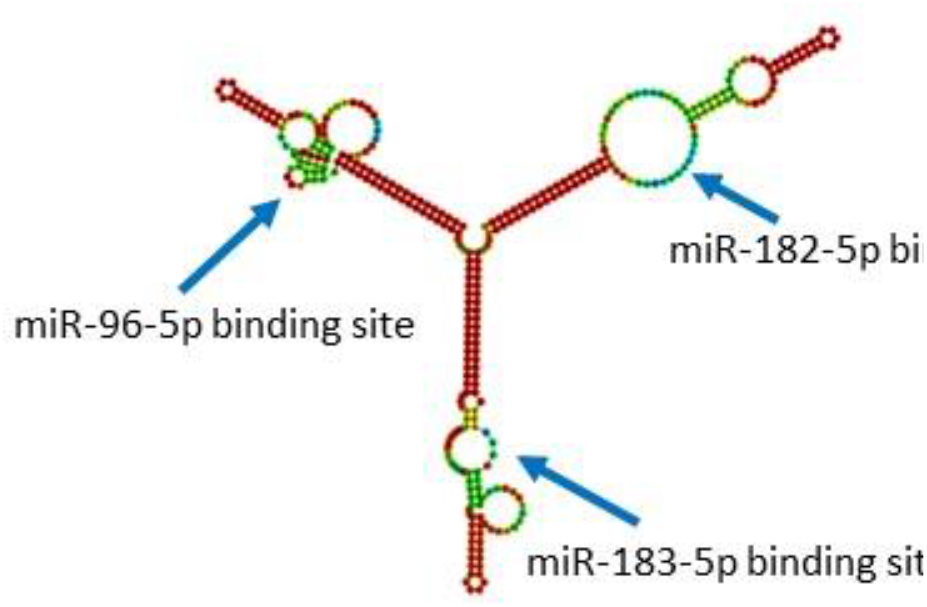
The structure of the TuD miR-183/96/182 transcript containing binding sites for the individual members of the miR-183/96/182 cluster. Picture generated by the RNAfold tool.

**Supplementary Figure 4:**
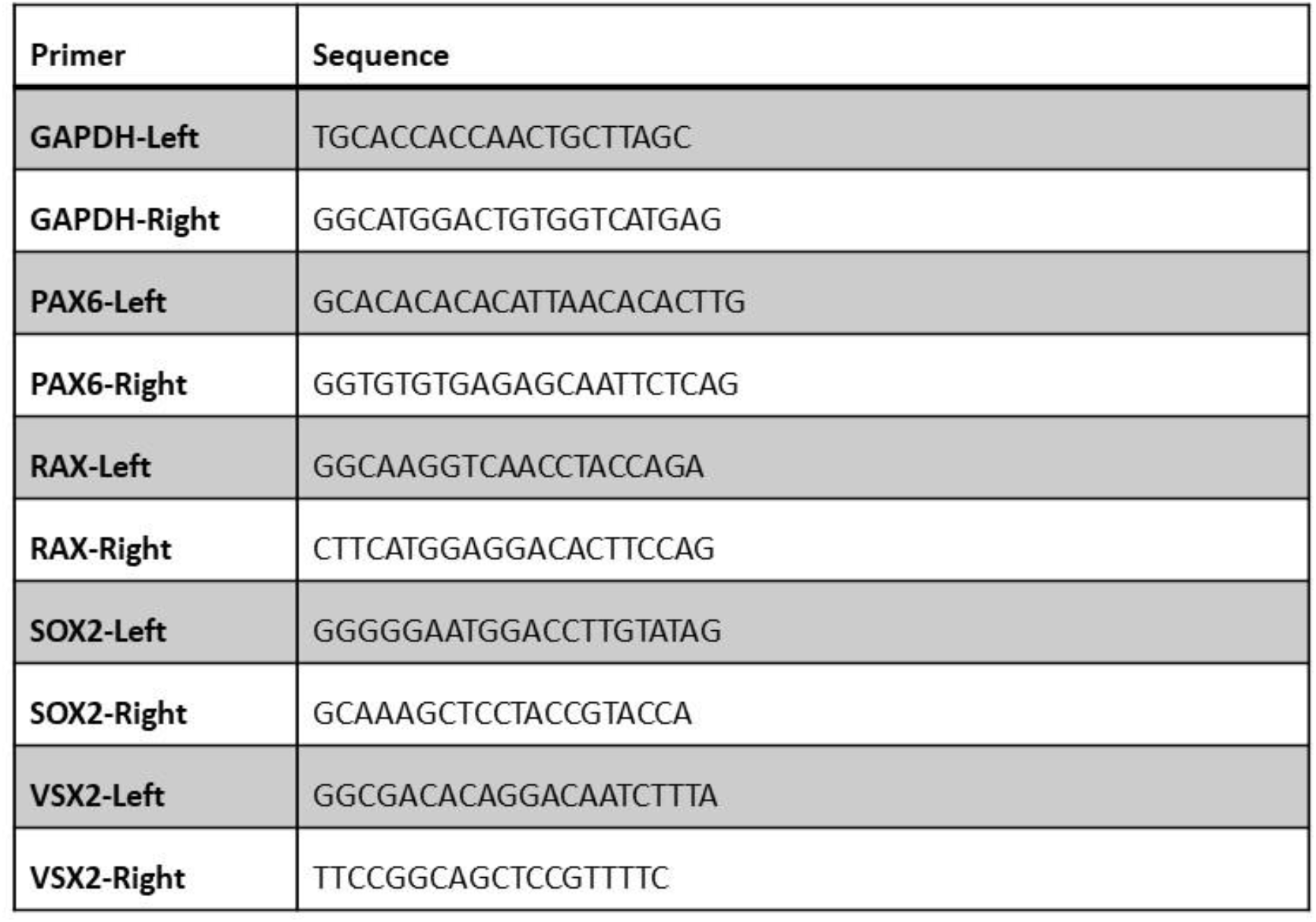
Primer sequences used in this study.

**Supplementary Figure 5:**
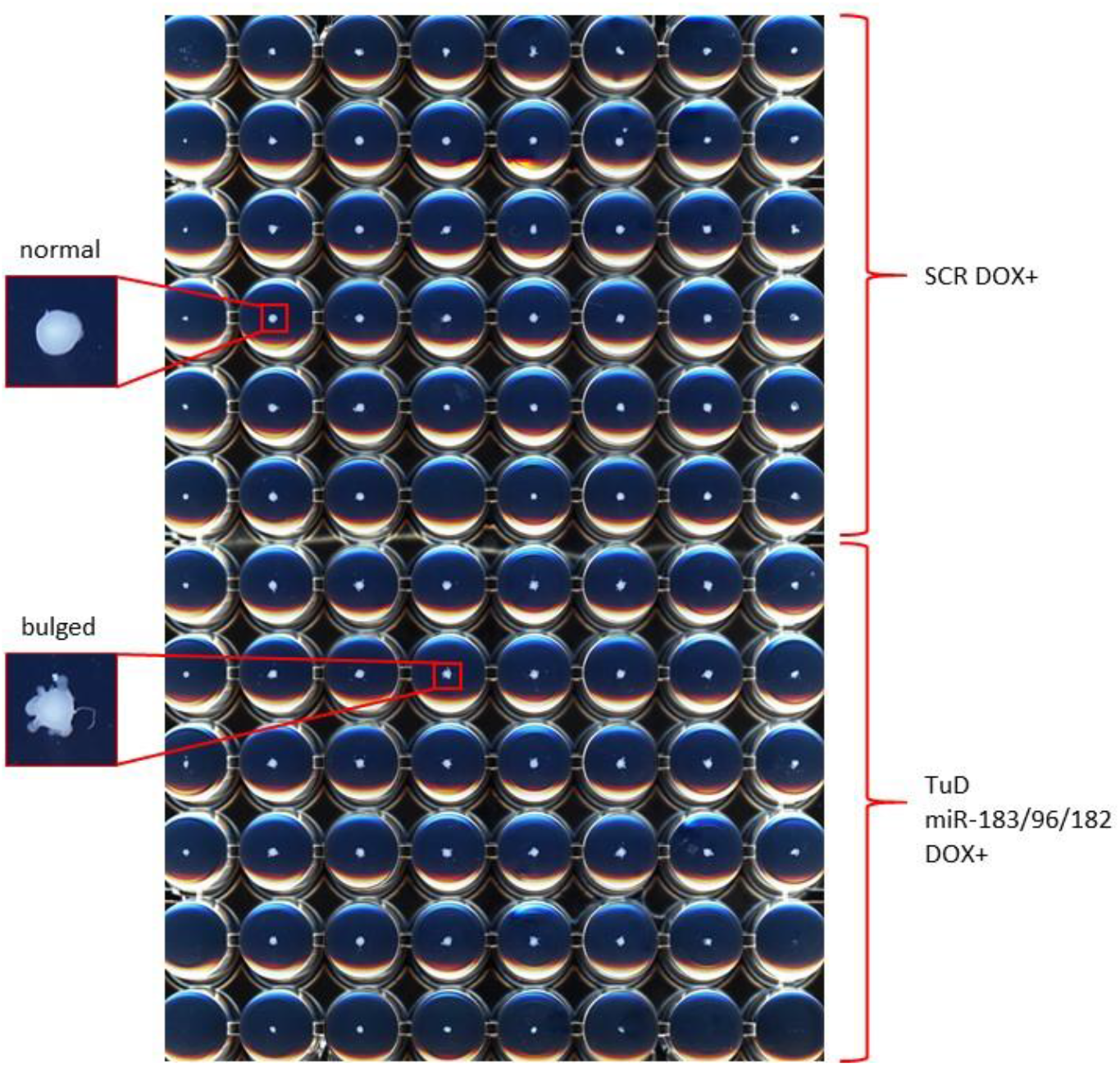
Representative picture of the scanned 96 well plate containing retinal organoids at D25 that was used for organoid circularity assessment.

## References

1 Dambal S, Shah M, Mihelich B, et al. The microRNA-183 cluster: the family that plays together stays together. Nucleic Acids Res 2015;43:7173–7188.

2 Hollensen AK, Bak RO, Haslund D, et al. Suppression of microRNAs by dual-targeting and clustered Tough Decoy inhibitors. RNA Biol 2013;10:406–414.

3 Barta T, Peskova L, Hampl A. miRNAsong: a web-based tool for generation and testing of miRNA sponge constructs in silico. Sci Rep 2016;6:36625.

4 Eshtad S, Mavajian Z, Rudd SG, et al. hMYH and hMTH1 cooperate for survival in mismatch repair defective T-cell acute lymphoblastic leukemia. Oncogenesis 2016;5:e275.

5 Peskova L, Cerna K, Oppelt J, et al. Oct4-mediated reprogramming induces embryonic-like microRNA expression signatures in human fibroblasts. Sci Rep 2019;9:15759.

6 Kuwahara A, Ozone C, Nakano T, et al. Generation of a ciliary margin-like stem cell niche from self-organizing human retinal tissue. Nat Commun 2015;6:6286.

7 Hallam D, Hilgen G, Dorgau B, et al. Human-Induced Pluripotent Stem Cells Generate Light Responsive Retinal Organoids with Variable and Nutrient-Dependent Efficiency. Stem Cells 2018;36:1535–1551.

8 Barta T, Peskova L, Collin J, et al. Brief Report: Inhibition of miR-145 Enhances Reprogramming of Human Dermal Fibroblasts to Induced Pluripotent Stem Cells. Stem Cells 2016;34:246–251.

9 Chou C-H, Shrestha S, Yang C-D, et al. miRTarBase update 2018: a resource for experimentally validated microRNA-target interactions. Nucleic Acids Res 2018;46:D296–D302.

10 Huang DW, Sherman BT, Lempicki RA. Systematic and integrative analysis of large gene lists using DAVID bioinformatics resources. Nat Protoc 2009;4:44–57.

11 Cline MS, Smoot M, Cerami E, et al. Integration of biological networks and gene expression data using Cytoscape. Nat Protoc 2007;2:2366–2382.

12 Shannon P, Markiel A, Ozier O, et al. Cytoscape: A Software Environment for Integrated Models of Biomolecular Interaction Networks. Genome Res 2003;13:2498–2504.

13 Susaki EA, Tainaka K, Perrin D, et al. Whole-brain imaging with single-cell resolution using chemical cocktails and computational analysis. Cell 2014;157:726–739.

14 Tainaka K, Kubota SI, Suyama TQ, et al. Whole-body imaging with single-cell resolution by tissue decolorization. Cell 2014;159:911–924.

15 Vejnar CE, Blum M, Zdobnov EM. miRmap web: Comprehensive microRNA target prediction online. Nucleic Acids Res 2013;41:W165–168.

16 Farzaneh M, Alishahi M, Derakhshan Z, et al. The Expression and Functional Roles of miRNAs in Embryonic and Lineage-Specific Stem Cells. Curr Stem Cell Res Ther 2019;14:278–289.

17 Giorgi Silveira R, Perelló Ferrúa C, do Amaral CC, et al. MicroRNAs expressed in neuronal differentiation and their associated pathways: systematic review and bioinformatics analysis. Brain Res Bull 2020.

18 Zeng Z-L, Lin X-L, Tan L-L, et al. MicroRNAs: Important Regulators of Induced Pluripotent Stem Cell Generation and Differentiation. Stem Cell Rev Rep 2018;14:71–81.

19 Ran X, Xiao C-H, Xiang G-M, et al. Regulation of Embryonic Stem Cell Self-Renewal and Differentiation by MicroRNAs. Cell Reprogram 2017;19:150–158.

20 Zuber ME, Gestri G, Viczian AS, et al. Specification of the vertebrate eye by a network of eye field transcription factors. Development 2003;130:5155–5167.

21 Chow RL, Altmann CR, Lang RA, et al. Pax6 induces ectopic eyes in a vertebrate. Development 1999;126:4213–4222.

22 Halder G, Callaerts P, Gehring WJ. Induction of ectopic eyes by targeted expression of the eyeless gene in Drosophila. Science 1995;267:1788-1792.

23 Czerny T, Halder G, Kloter U, et al. twin of eyeless, a Second Pax-6 Gene of Drosophila, Acts Upstream of eyeless in the Control of Eye Development. Molecular Cell 1999;3:297–307.

24 Manuel M, Pratt T, Liu M, et al. Overexpression of Pax6 results in microphthalmia, retinal dysplasia and defective retinal ganglion cell axon guidance. BMC Developmental Biology 2008;8:59.

25 Du Z-W, Ma L-X, Phillips C, et al. miR-200 and miR-96 families repress neural induction from human embryonic stem cells. Development 2013;140:2611–2618.

26 Xu S, Witmer PD, Lumayag S, et al. MicroRNA (miRNA) transcriptome of mouse retina and identification of a sensory organ-specific miRNA cluster. J Biol Chem 2007;282:25053–25066.

27 Xu S. microRNA expression in the eyes and their significance in relation to functions. Progress in Retinal and Eye Research 2009;28:87–116.

28 Oron-Karni V, Farhy C, Elgart M, et al. Dual requirement for Pax6 in retinal progenitor cells. Development 2008;135:4037–4047.

29 Lumayag S, Haldin CE, Corbett NJ, et al. Inactivation of the microRNA-183/96/182 cluster results in syndromic retinal degeneration. Proc Natl Acad Sci USA 2013;110:E507–516.

30 Busskamp V, Krol J, Nelidova D, et al. miRNAs 182 and 183 are necessary to maintain adult cone photoreceptor outer segments and visual function. Neuron 2014;83:586–600.

